# Cortical tracking of unheard formant modulations derived from silently presented lip movements and its decline with age

**DOI:** 10.1101/2021.04.13.439628

**Authors:** Nina Suess, Anne Hauswald, Patrick Reisinger, Sebastian Rösch, Anne Keitel, Nathan Weisz

**Author notes:** Address for correspondence: Nina Suess, University of Salzburg, Centre for Cognitive Neuroscience, Hellbrunner Straße 34, A-5020 Salzburg, Austria/Europe. shared senior authors (Anne Keitel, Nathan Weisz).

## Abstract

The integration of visual and auditory cues is crucial for successful processing of speech, especially under adverse conditions. Recent reports have shown that when participants watch muted videos of speakers, the phonological information about the acoustic speech envelope is tracked by the visual cortex. However, the speech signal also carries much richer acoustic details, e.g. about the fundamental frequency and the resonant frequencies, whose visuo-phonological transformation could aid speech processing. Here, we investigated the neural basis of the visuo-phonological transformation processes of these more fine-grained acoustic details and assessed how they change with ageing. We recorded whole-head magnetoencephalography (MEG) data while participants watched silent intelligible and unintelligible videos of a speaker. We found that the visual cortex is able to track the unheard intelligible modulations of resonant frequencies and the pitch linked to lip movements. Importantly, only the processing of intelligible unheard formants decreases significantly with age in the visual and also in the cingulate cortex. This is not the case for the processing of the unheard speech envelope, the fundamental frequency or the purely visual information carried by lip movements. These results show that unheard spectral fine-details (along with the unheard acoustic envelope) are transformed from a mere visual to a phonological representation. Aging affects especially the ability to derive spectral dynamics at formant frequencies. Since listening in noisy environments should capitalize on the ability to track spectral fine-details, our results provide a novel focus on compensatory processes in such challenging situations.

## 1 Introduction

Speech understanding is a multisensory process that requires diverse modalities to work together for an optimal experience. Congruent audiovisual input is especially crucial for understanding speech in noise (Crosse et al., 2016; Sumby & Pollack, 1954), highlighting the importance of visual cues in speech processing studies. One hypothesis is that activation from visual speech directly modulates activation in auditory cortex, although the results have been mixed and a lot of questions remain unanswered (Bernstein & Liebenthal, 2014; Keitel et al., 2020). One important question regards the nature of the representation in the visual cortex, and whether it is strictly visual or already tracks acoustic information that is associated with the visual input (for non-speech stimuli see e.g. Escoffier et al., 2015). A first approach to address this showed that occipital activation elicited by silent lip reading also reflects dynamics of the acoustic envelope (O’Sullivan et al., 2017). Further evidence that the visual cortex is able to track certain aspects of speech by visual cues alone comes from a recent study by Hauswald et al. (2018). Evidently, it has been shown that visual speech contributes substantially to audiovisual speech processing in the sense that the visual cortex is able to extract phonological information from silent lip movements in the theta-band (4-7 Hz). Crucially, this tracking is dependent on the intelligibility of the silent speech, with absent tracking when the silent visual speech is unintelligible. Another study supports the former findings and extends the present framework by providing evidence that the visual cortex passes information to the angular gyrus, which extracts slow features (below 1 Hz) from lip movements, which are then mapped onto auditory features and passed on to auditory cortices for better speech comprehension (Bourguignon et al., 2020). These findings underline the importance of slow frequency properties of visual speech for enhanced speech comprehension from both the delta (0.5-3 Hz) and theta-band (4-7 Hz), especially due to frequencies between 1-7 Hz being crucial for comprehension (Giraud & Poeppel, 2012). Moreover, the spectral profile of lip movements is also settled within this range (Park et al., 2016).

Recent behavioural evidence describes that spectral fine details can also be extracted by observation of lip movements (Plass et al., 2020). This raises the interesting question whether this information is also represented at the level of the visual cortex, analogous to the envelope as shown previously (Hauswald et al., 2018). Particularly relevant spectral fine details are formant peaks around 2500 Hz, which are indicated to be modulated in the front cavity (Badin et al., 1990). This corresponds to expansion and contraction of the lips (Plass et al., 2020), thus having a relationship with certain lip movements and could therefore be extracted for important phonological cues.

Furthermore, not only resonant frequencies, but also the fundamental frequency (or pitch contour) plays an important role in speech understanding in noisy environments (Hopkins et al., 2008), and could potentially be extracted from silent lip movements. Whether the visual cortex is able to track formant and pitch information in (silent) visual speech, has not been investigated to date.

Knowledge on how the brain is processing speech is also vital when it comes to ageing, potentially with regards to age-related hearing loss (Liberman, 2017). Several studies have investigated the influence of age on speech comprehension, with results that signify ageing is, in most cases, accompanied by listening difficulties, especially in noise (Tun & Wingfield, 1999; Wong et al., 2009). Furthermore, while the auditory tracking of a speech-paced stimulation (~ 3 Hz) is less consistent in older adults compared to younger adults, alpha oscillations are enhanced in younger adults during attentive listening, suggesting declined top-down inhibitory processes that support selective suppression of irrelevant information (Henry et al., 2017). Older adults also indicate a compensatory mechanism when processing degraded speech especially in anterior cingulate cortex (ACC) and middle frontal gyrus (Erb & Obleser, 2013). Additionally, the temporal processing of auditory information is altered in the ageing brain, pointing to decreased selectivity for temporal modulations in primary auditory areas (Erb et al., 2020). Those studies reinforce a distinctive age-related alteration in processing auditory speech. This raises the question whether we also see an impact of age on audiovisual speech processing, an issue that has not been addressed so far.

Combining the important topics mentioned above, this study aims to answer two critical questions regarding audiovisual speech processing: First, we ask if the postulated visuo-phonological transformation process in visual cortex mainly represents global energy modulations (i.e. speech envelope) or if it also entails spectral fine details (like formant or pitch curves). Second, we question if visuo-phonological transformation is subject to age-related decline. To the best of our knowledge, this study presents first neurophysiological evidence that the visual cortex is not only able to extract the unheard speech envelope, but also unheard formant and pitch information from lip movements. Crucially, we observed an age-related decline that mainly affects tracking of the formants (and to some extent the envelope and the fundamental frequency). Interestingly, we observed different tracking properties for different brain regions and frequencies: While tracking intelligible formants declines reliably in occipital and cingulate cortex for both delta and theta, we observed a decline of theta-tracking just in occipital cortex, suggesting different age-related effects in different brain regions. Our results suggest that the ageing brain deteriorates in deriving spectral fine-details linked to the visual input, a process that could contribute to perceptual difficulties in challenging listening situations.

## 2 Materials and methods

### 2.1 Participants

We recruited 50 participants (28 females; 2 left-handed; mean age: 37.96 years; SD: 13.33 years, range: 19-63 years) for the experiment. All participants had normal or corrected-to-normal eyesight, self-reported normal hearing and no neurological disorders. All participants received either a reimbursement of €10 per hour or course credits for their participation. All participants signed an informed consent form. The experimental procedure was approved by the Ethics Committee of the University of Salzburg.

### 2.2 Stimuli

Videos were recorded with a digital camera (Sony NEX FS100) at a rate of 50 frames per second, the corresponding audio files were recorded at a sampling rate of 48 kHz. The videos were spoken by two female native German speakers. The stimuli were taken from the book “Das Wunder von Bern” (“The Miracle of Bern”; https://www.aktion-mensch.de/inklusion/bildung/bestellservice/materialsuche/detail?id=62) which was provided in an easy language. The easy language does not include any foreign words, has a coherent verbal structure and is facile to understand. We used simple language to avoid that limited linguistic knowledge is interfering with possible lip reading abilities. 24 pieces of text were chosen from the book and recorded from each speaker, lasting between 33 and 62 seconds, thus resulting in 24 videos. Additionally, all videos were reversed, which resulted in 24 forward videos and 24 corresponding backward videos. Forward and backward audio files were extracted from the videos and used for the data analysis. Half of the videos were randomly selected to be presented forward and the remaining half to be presented backward. The videos were back-projected on a translucent screen in the centre of the screen by a Propixx DLP projector (VPixx technologies, Canada) with a refresh rate of 120 Hz per second and a screen resolution of 1920 × 1080 pixels. The translucent screen was placed ~110 cm in front of the participant and had a screen diagonal of 74 cm. One speaker was randomly chosen per subject and kept throughout the experiment, so each participant only saw one speaker.

### 2.3 Procedure

Participants were first instructed to take part in an online study, in which their behavioural lip reading abilities were tested, and in which they were asked about their subjective hearing impairment. This German lip reading test is available as SaLT (Salzburg Lipreading Test) (Suess et al., 2021). Participants were presented with silent videos of numbers, words and sentences and could watch every video twice. They then had to write down the words they thought they had understood from the lip movements. This online test lasted approximately 40 minutes and could be conducted at home or right before the experiment in the MEG-lab. After completing the behavioural experiment, the MEG experiment started. Participants were instructed to pay attention to the lip movements of the speakers and passively watch the mute videos. They were presented with 6 blocks of videos, and in each block, 2 forward and 2 backward videos were presented in a random order. The experiment lasted about an hour including preparation. The experimental procedure was programmed in Matlab with the Psychtoolbox-3 (Brainard, 1997) and an additional class-based abstraction layer (https://gitlab.com/thht/o_ptb) programmed on top of the Psychtoolbox (Hartmann & Weisz, 2020).

### 2.4 Data acquisition

Brain activity was measured using a 306-channel whole head MEG system with 204 planar gradiometers and 102 magnetometers (Neuromag TRIUX, Elekta), a sampling rate of 1000 Hz and an online highpass-filter of 0.1 Hz. Before entering the magnetically shielded room (AK3B, Vakuumschmelze, Hanau, Germany), the head shape of each participant was acquired using approximately 500 digitized points on the scalp, including fiducials (nasion, left and right pre-auricular points) with a Polhemus Fastrak system (Polhemus, Vermont, USA). The head position of each individual participant relative to the MEG sensors was controlled once before each experimental block. Vertical and horizontal eye movements and electrocardiographic data was also recorded, but not used for preprocessing. The continuous MEG data was then preprocessed off-line with the signal space separation method from the Maxfilter software (Elekta Oy, Helsinki, Finland) to correct for different head positions across blocks and to suppress external interference (Taulu et al., 2005).

### 2.5 Data analysis

#### 2.5.1 Preprocessing

Acquired datasets were analysed using the Fieldtrip toolbox (Oostenveld et al., 2011). The maxfiltered MEG data were highpass-filtered at 1 Hz using a finite impulse response (FIR) filter (Kaiser window, order 440). For extracting physiological artefacts from the data, 60 principal components were calculated. Via visual inspection, the components displaying eye movements, heartbeat and external power noise from the nearby train tracks (16.67 Hz) were removed from the data. We removed on average 2.24 components per participant (*SD* = 0.65). The data were then lowpass-filtered at 30 Hz and corrected for the delay between the stimulus computer and the screen inside the chamber (9 ms for each video). We then resampled the data to 150 Hz and segmented them in 2-second trials to increase the signal-to-noise ratio.

#### 2.5.2 Source projection of MEG data

We used either a standard structural brain from the Montreal Neurological Institute (MNI, Montreal, Canada) or, where possible, the individual structural MRI (20 participants) and warped it to match the individual’s fiducials and head shape as accurately as possible (Mattout et al., 2007). A 3-D grid with 1-cm resolution and 2982 voxels based on an MNI template brain was morphed into the brain volume of each participant. This allows group-level averaging and statistical analysis as all the grid points in the warped grid belong to the same brain region across subjects. These aligned brain volumes were also used for computing single-shell head models and leadfields (Nolte, 2003). By using the leadfields and the common covariance matrix (pooling data from all blocks), a common LCMV beamformer spatial filter was computed (Veen et al., 1997).

#### 2.5.3 Extraction of lip area, acoustic speech envelope, formants and pitch

The lip area of the visual speech was extracted using a MATLAB script adapted from Park et al. (2016). This data was then upsampled to 150 Hz to match the downsampled preprocessed MEG signal. The acoustic speech envelope was extracted with the Chimera toolbox from the audio files corresponding to the videos which constructs nine frequency bands in the range of 100-10000 Hz as equidistant on the cochlear map (Smith et al., 2002). Then the sound stimuli were band-pass filtered in these bands with a 4th-order Butterworth filter to avoid edge artefacts. For each of the frequency bands, the envelopes were calculated as absolute values of the Hilbert transform and then averaged to get the full-band envelope for coherence analysis (Gross et al., 2013; Keitel et al., 2017). This envelope was then downsampled to 150 Hz to match the preprocessed MEG signal. The resonant frequencies (or formants) were extracted using the Burg method implemented in Praat 6.0.48 (Boersma & Weenink, 2019). Up to 5 formants were extracted from each audio file to make sure that the relevant formants were extracted thoroughly. For analysis purposes, just F2 and F3 were averaged and used. Those two formants fluctuate around 2500 Hz and tend to merge into a single peak when pronouncing certain consonant-vowel combinations (Badin et al., 1990). The mentioned merging process is taking place in the front region of the oral cavity and can therefore also be seen by observing lip movements (Plass et al., 2020). The formants were extracted at a rate of 200 Hz for the sake of simplicity and then downsampled to 150 Hz. The pitch (or fundamental frequency, f0) was extracted using the Matlab Audio Toolbox function *pitch.m* with default options (extraction between 50 and 400 Hz) at a rate of 100 Hz and then upsampled to 150 Hz.

#### 2.5.4 Coherence calculation

We calculated the cross-spectral density between the lip area, the unheard acoustic speech envelope, the averaged F2 and F3 formants and the pitch and every virtual sensor with a multi-taper frequency transformation (1-25 Hz in 0.5 Hz steps, 3 Hz smoothing). Then we calculated the coherence between the activity of every virtual sensor and the lip area, the acoustic speech envelope, the averaged formant curve of F2 and F3 and the pitch curve, which we will refer to as lip-brain coherence, envelope-brain coherence, formant-brain coherence and pitch-brain coherence, respectively, in the manuscript.

### 2.6 Statistical analysis

To test for differences in source space in occipital cortex for forward and backward coherence values, we extracted all voxels labeled as “occipital cortex” in the Automated Anatomical Labeling (AAL) atlas (Tzourio-Mazoyer et al., 2002) for a predefined region-of-interest analysis (Hauswald et al., 2018). We then contrasted forward and backward conditions using two-tailed dependent-samples *t*-tests on the averaged coherence values for the frequency bands of interest (1-7 Hz). This was done separately for the lip-brain coherence, the envelope-brain coherence, the formant-brain coherence and the pitch-brain coherence. In a first step, we decided to average over the delta (1-3 Hz) and theta (4-7 Hz) frequency bands since they carry important information in general on speech processing (phrasal and syllabic processing, respectively) (Giraud & Poeppel, 2012). Moreover, previous studies investigated lip movement related activity either in the delta-band (Bourguignon et al., 2020; Park et al., 2016) or the theta-band (Hauswald et al., 2018), leading us to also do a follow-up analysis separately for the different frequency bands (described later in this section).

To generate a normalized contrast between processing of forward (intelligible) and backward (unintelligible) lip movements, we subtracted the backward coherence values from the forward coherence values for our respective measures (lip-brain coherence, unheard speech envelope-brain coherence, unheard formant-brain coherence and unheard pitch-brain coherence). From now on, we refer to this normalized contrast as “Intelligibility index”, which quantifies the differences in coherence between intelligible and unintelligible visual speech. For testing the relationship between the four different intelligibility indices (lip-brain, envelope-brain, formant-brain and pitch-brain) and age, we conducted a voxelwise correlation with age. To control for multiple comparisons, we used a non-parametric cluster-based permutation test (Maris & Oostenveld, 2007). Here, clusters of correlation coefficients being significantly different from zero (showing p-values < 0.05) were identified and their respective *t*-values were extracted and summed up to get a cluster-level test statistic. Random permutations of the data were then drawn by reordering the behavioural data (in our case age) across participants. After each permutation, the maximum cluster level *t*-value was recorded, generating a reference distribution of cluster-level *t*-values (using a Monte Carlo procedure with 1000 permutations). Cluster *p*-values were estimated as the proportion of cluster *t*-values in the reference distribution exceeding the cluster *t*-values observed in the actual data. Significant voxels (which were only found in the correlation between the formant-brain index and age) were then extracted and averaged for data-driven ROIs (occipital cortex and cingulate cortex) which were defined using the Automated Anatomical Labeling (AAL) atlas (Tzourio-Mazoyer et al., 2002). These data-driven ROIs were then applied to all intelligibility indices to make the ROI analysis comparable. We then fitted four linear models using the function *lm* from the stats package in R to investigate if age could predict the change in the calculated intelligibility indices and to visualize the statistical effects of the whole brain analysis. To further clarify the relationship between age and the processing of intelligible and unintelligible lip movements and to unravel the dynamics in our whole brain correlation analysis, we split our participants into two groups by the median (young: people < 37, N=25, older: people > 37, N=25). We then calculated a repeated-measures ANOVA with 2 conditions: age (young vs. older) and intelligibility (forward vs. backward visual speech) for our data-driven ROIs separately (occipital cortex and cingulate cortex) using the *stats* package in R. To further investigate the effects between age and intelligibility, we conducted post-hoc tests with Bonferroni correction using the function *PostHocTest*. The last step consisted of a follow-up analysis where we decided to separate the averaged frequency-bands (delta and theta) again to unravel possible differences of our effect dependent on the frequency-band. We again conducted a voxelwise correlation with age separately for the delta-band (1-3 Hz) and for the theta-band (4-7 Hz) with the already described non-parametric cluster-based permutation test for all described intelligibility indices. Finally, we extracted the values from the voxel with the lowest *t*-value (for the delta and theta-band, respectively) and fitted a linear model again to investigate if age could predict the change in the intelligibility indices and to visualize the statistical effects of the whole brain analysis.

## 3 Results

### 3.1 Behavioural results

We investigated participants’ lip reading abilities in a separate experiment that was conducted before the MEG session. They were presented with silent videos of spoken numbers, words, and sentences, and the task was to write down what they had understood just from the lip movements alone. A detailed description of the behavioural task will be published in a separate paper (Suess et al., 2021). 43 of the 50 participants completed the behavioural experiment. 4 people had to be excluded because there were problems with the data acquisition and their answers were not saved. While the recognition rate for the numbers were high (M = 60.83%, SE = 2.93%), lip reading abilities for complex stimuli (words and sentences) were low in general (words: M = 30.57%, SE = 1.82%; sentences: M = 8.83%, SE = 1.26%). Participants had an average total score of 33.41% (SE = 1.75%, Figure 2A). Investigating if age could predict the total test score revealed that those two variables were uncorrelated (F(1, 37) = .191, p = .664, R^2^ = −0.021), Figure 2B), showing that in our sample, behavioural lip reading abilities are not changing with age. This is consistent with our study on general lip reading abilities in the German language (Suess et al., 2021), but different to other studies indicating higher lip reading abilities in younger individuals (Feld & Sommers, 2009; Tye-Murray et al., 2007b). Participants also completed a questionnaire on subjective hearing impairment (APHAB, Löhler et al., (2014)). Further investigating the relationship between subjective hearing impairment and test score also revealed no significant effect (F(1, 37) = .104, p = .75, R^2^ = −0.024) in the current sample. This is in line with studies investigating hearing impairment in older adults (Tye-Murray et al., 2007a), but not supporting our own results which show a relationship between self-reported hearing impairment and lip reading abilities (Suess et al., 2021). However, as the current study was aiming to test normal hearing individuals with restricted variance in hearing impairment, those results cannot be compared directly to Suess et al. (2021), which also included individuals with severe hearing loss as well as prelingually deaf individuals.

**Figure 1:**
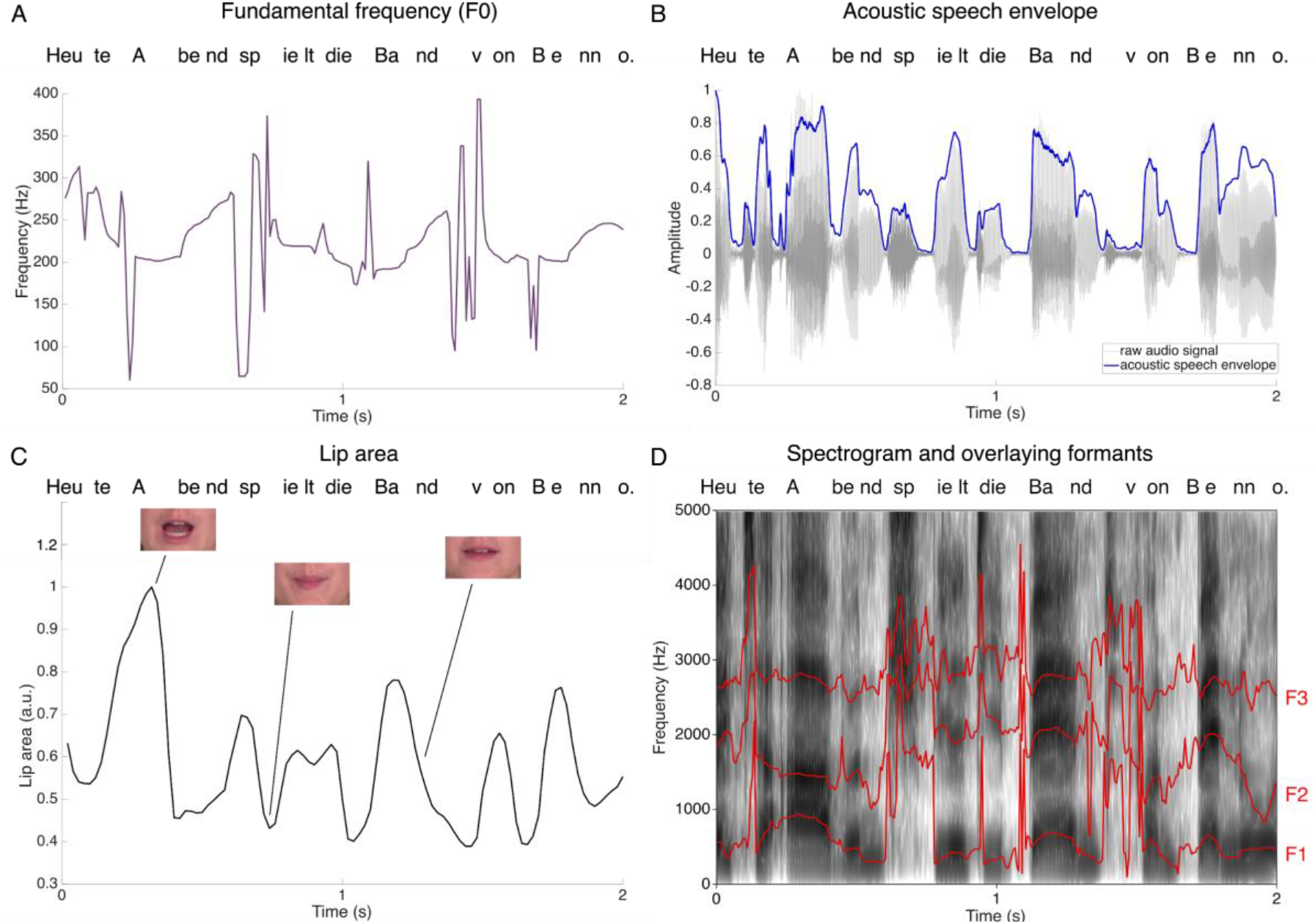
Example time series for a 2 second forward section of all the parameters used for coherence calculation. A) Example time series of the fundamental frequency extracted with the pitch.m MATLAB function. B) Example audio signal and the acoustic speech envelope (in blue). C) Lip area extracted from the video frames with the MATLAB script adapted from Park et al. (2016). D) Example spectrogram with overlaying formants (F1-F3, red lines) extracted with Praat.

**Figure 2:**
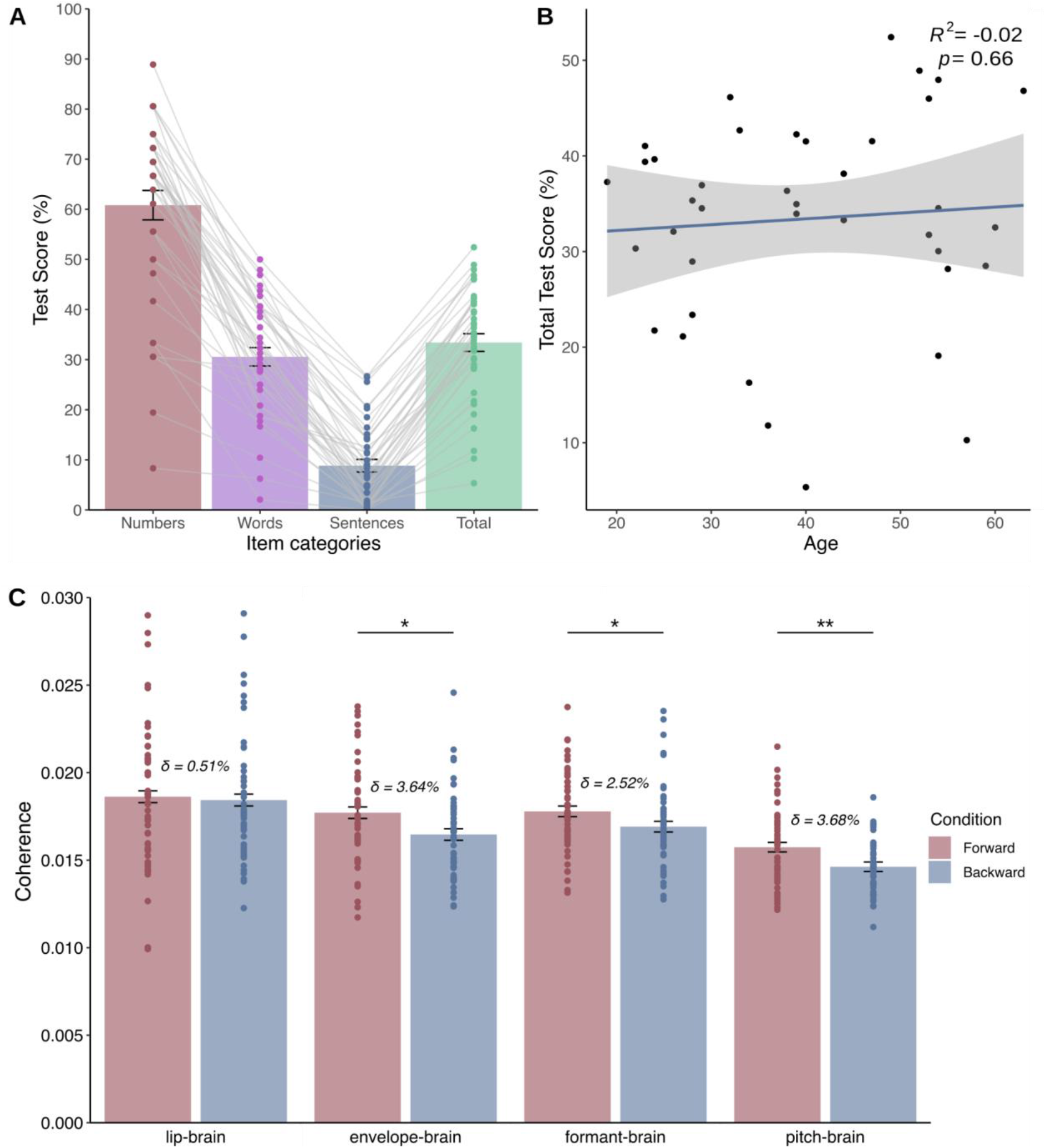
Behavioural data and comparison of information tracking in visual cortex. A) Behavioural lip reading abilities. Participants recognized numbers the most, followed by words and sentences. B) Correlation between age and total test score revealed no significant correlation (p = 0.66), suggesting that lip reading abilities do not change with age. Blue line depicts regression line, shaded areas depict standard error of mean (SE). C) Mean values extracted from all voxels in occipital cortex showing no significant differences in lip-brain coherence (p = 0.694), but showing significant differences in unheard envelope-brain coherence (p = 0.01), formant-brain coherence (p = 0.047) and unheard pitch-brain coherence (p = 0.005) between forward and backward presentation of visual speech. Error bars represent 1 standard error of mean for within-subject designs (O’Brien & Cousineau, 2014), δ indicates the relative change between forward and backward conditions in percent.

### 3.2 Visuo-phonological transformation is carried by both tracking of global envelope and spectral fine-details during presentation of intelligible silent lip movements

We calculated the coherence between the MEG data and the lip envelope, the unheard acoustic speech envelope, the unheard resonant frequencies and the unheard pitch (from now on called lip-brain coherence, envelope-brain coherence, formant-brain coherence, and pitch-brain coherence, respectively). As the visuo-phonological transformation process is likely taking place in visual areas (Hauswald et al., 2018), we defined the occipital cortex using the AAL atlas (Tzourio-Mazoyer et al., 2002) as a predefined region-of-interest and averaged over all voxels from this ROI. We then compared the mean for the coherence of the presented forward videos (intelligible lip movements) with the mean of the presented backward videos (unintelligible lip movements) separately for the lip-brain coherence, the envelope-brain coherence, the formant-brain coherence and the pitch-brain coherence. While there was no significant difference in lip-brain coherence for intelligible and unintelligible visual speech (*t*(49) = 0.396, *p* = 0.694, *d* = 0.056), we found a significant difference in unheard envelope-brain coherence for intelligible and unintelligible visual speech (*t*(49) = 2.679, *p* = 0.01, *d* = 0.379). Most importantly, we found a significant difference also for the unheard formant-brain coherence (t(49) = 2.039, *p* = 0.047, *d* = 0.288) and for the unheard pitch-brain coherence for intelligible and unintelligible visual speech (t(49) = 2.91, *p* = 0.005, *d* = 0.411, all in Figure 2C). The results on the tracking of lip movements are in line with former findings, showing that the visual cortex tracks these regardless of intelligibility, but point to different tracking properties dependent on the intelligibility of the unheard speech envelope. Interestingly, we show here that the visual cortex is also able to distinguish between unheard intelligible and unintelligible formants (or resonant frequencies) and pitch (or F0) modulations extracted from the spectrogram, showing that also unheard intelligible spectral details are extracted from visual speech and represented at the level of the visual cortex.

#### 3.3 Spectral fine-detail tracking rather than global envelope tracking is altered in the ageing population

We were then interested in how the visuo-phonological transformation process is influenced by age. So we calculated a voxelwise correlation between the intelligibility index (difference between coherence for forward videos and coherence for backward videos) separately for our coherence indices (lip-brain, envelope-brain, formant-brain and pitch-brain) and the age of the participants. We neither found a significant correlation between the intelligibility index of the lip-brain coherence and age (*p* = 1, cluster-corrected) nor between the intelligibility index of the unheard envelope-brain coherence and age (*p* = 0.09, cluster-corrected). Also, the correlation between the intelligibility index of the unheard pitch-brain coherence was statistically not significant (p = 0.07, cluster-corrected). However, the overall trend for the envelope-brain and the pitch-brain coherence was to decline with age. Interestingly, we did find a significant negative correlation between the intelligibility index of the unheard formant-brain coherence and age (*p* = 0.002, cluster-corrected), strongest in occipital cortex and cingulate cortex (lowest *t*-value: −4.124, MNI [40 −90 0], Figure 3A). To further investigate the effects, we extracted the voxels showing a statistical effect in our whole brain analysis (Figure 3A) and divided them into occipital voxels and voxels from the cingulate cortex using the AAL atlas (Tzourio-Mazoyer et al., 2002).

**Figure 3:**
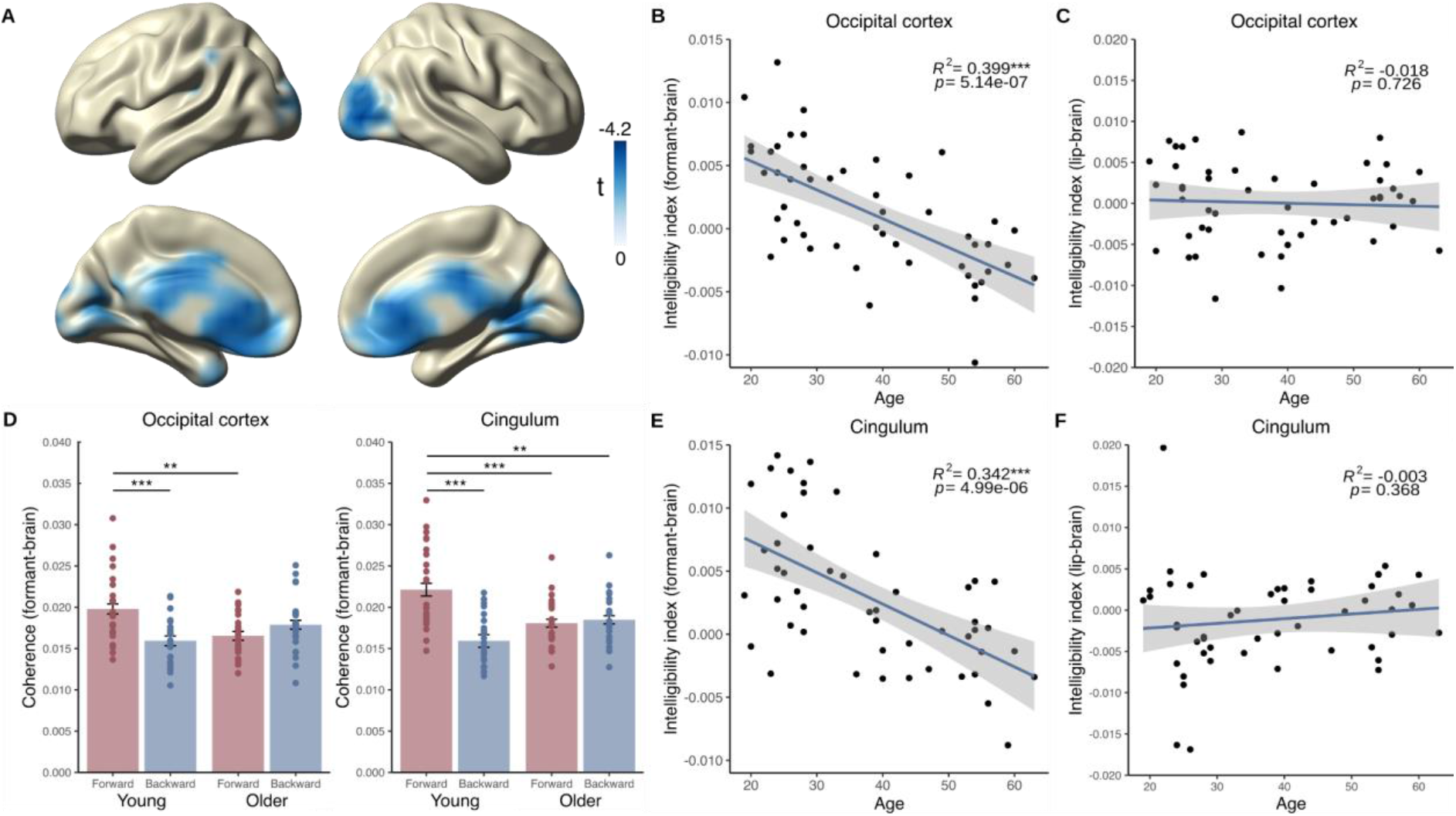
Correlation between age and intelligibility index (i.e. difference in forward vs. backward tracking) and comparison of age-groups. A) Statistical values of the voxelwise correlation of the intelligibility index (forward formant-brain coherence - backward formant-brain coherence) with age (averaged over 1-7 Hz, p < 0.05, cluster-corrected) showing a strong decrease of intelligibility tracking in occipital regions and in cingulate cortex. B) Correlation of formant-brain intelligibility index in significant occipital voxels extracted from A showing a significant correlation with age (p = 5.14e-07). C) Correlation of lip-brain intelligibility index in significant occipital voxels extracted from A showing a not significant correlation with age (p = 0.726). D) Formant-brain coherence separated for age and for forward and backward presented visual speech for different ROIs. Coherence values from occipital cortex indicating significant differences between forward and backward tracking in the young group (p = 0.0004), but not in the older group (p = 0.467), and also a difference between forward tracking in the young group and forward tracking in the older group (p = 0.004). Coherence values from cingulum indicating significant differences between forward and backward tracking in the young group (p = 1.1e-07), but not in the older group (p = 1.000), and also a difference between forward tracking in the young group and forward tracking in the older group (p = 0.0005). Additional significant effects were observed between the forward tracking in the young group and the backward tracking in the older group (p = 0.002). E) Correlation of formant-brain intelligibility index in significant voxels from cingulate cortex extracted from A showing a significant correlation with age (p = 4.99e-06). F) Correlation of lip-brain intelligibility index in significant voxels from cingulate cortex extracted from A showing a not significant correlation with age (p = 0.368). Blue lines depict regression lines, shaded areas depict standard error of mean (SE).

To investigate how strong the relationship between age and the different intelligibility indices is in our ROIs, we fitted four separate linear models. We started with the lip-brain index to exclude the possibility that our effect is due to visual processing. We found that age could not predict the lip-brain intelligibility index in any of the chosen ROIs (occipital cortex: F(1, 48) = 0.124, *p* = 0.727, *η^2^*= 0.002, Figure 3C; cingulate cortex: F(1, 48) = 0.825, *p* = 0.368, *η^2^*= 0.017, Figure 3F). On the contrary, we found that age could significantly predict the decrease in the formant-brain intelligibility index in both occipital areas (F(1, 48) = 33.59, *p* = 5.14e-07, η^2^= 0.412, Figure 3B) and cingulate cortex (F(1, 48) = 26.42, *p* = 4.99e-06, *η^2^*= 0.355, Figure 3E), suggesting an altered tracking process for the formants in ageing. Further fitting linear models to investigate the effects in our ROIs for the envelope-brain coherence and the pitch-brain coherence revealed that age could not significantly predict the envelope-brain index in occipital (F(1, 48) = 1.638, *p* = 0.207, *η^2^ =* 0.033) or cingulate cortex (F(1, 48) = 0.681, *p* = 0.413, *η^2^* = 0.014) and also not the pitch-brain index in occipital cortex (F(1, 48) = 2.584, *p* = 0.114, *η^2^ =* 0.051). However, age could significantly predict the pitch-brain index in cingulate cortex (F(1, 48) = 6.972, *p* = 0.011, *η^2^ =* 0.127). The lack of tracking differences between intelligible and unintelligible lip movements suggests that the visual cortex processes basic visual properties of lip movements, but that there are differential processing strategies for acoustic information associated with these lip movements. These results also suggest that processing of the pitch (or fundamental frequency) is altered to some extent in the ageing population, at least in cingulate cortex. In summary, the correlation between the envelope-brain index and age and the pitch-brain index and age seem to show a tendency in line with the relationship between the formant-brain index and age in the whole brain analysis. We see that effect sizes are biggest for the formant-brain index (occipital *η^2^ =* 0.412, cingulate *η^2^* = 0.355), followed by the pitch-brain index (occipital *η^2^* = 0.051, cingulate *η^2^* = 0.127). Lower effect sizes are found for the envelope-brain index (occipital *η^2^* = 0.033, cingulate *η^2^ =* 0.014) and the lip-brain index (occipital *η^2^ =* 0.002, cingulate *η^2^ =* 0.017) after extracting voxels from the data-driven ROI, adding to the evidence of a differential processing of speech properties in age.

### 3.4 Intelligibility effects are mainly carried by young individuals

To unravel the effects explained in sections 3.3, we reassessed the coherence values separately for forward and backward speech with respect to the age of our participants. Thus, we decided to split our sample into two age groups (younger vs. older) and calculated a 2×2 ANOVA on the averaged voxels that we extracted from figure 3A for the former calculated formant-brain coherence (for forward and backward coherence, respectively). We again separated them into two ROIs (occipital cortex and cingulate cortex) and calculated for each an ANOVA with the factors age (young vs. older) and intelligibility (forward formant-brain coherence vs. backward formant-brain coherence). We did not find a main effect of age in occipital cortex (F(1, 49) = 0.981, *p* = 0.324), but a main effect close to significance threshold of intelligibility (F(1, 49) = 3.627, *p =* 0.059). We also found a distinct interaction effect between age and intelligibility (F(1, 49) = 15.723, *p =* 0.0001, Figure 3D, occipital cortex). To further investigate the interaction effect, we calculated a post-hoc test with Bonferroni correction, which revealed a significant difference between the forward and backward conditions in the young group (*p =* 0.0004), but not in the older group (*p = 0.467*). Furthermore, we discovered a significant difference between the forward condition in the young group and the forward condition in the older group (*p* = 0.004), exhibiting that the young group is able to track the forward speech stronger than the older group. In cingulate cortex, we also did not find a main effect of age (F(1, 49) = 1.399, *p* = 0.239), but here we found a main effect of intelligibility (F(1, 49) = 16.474, *p* = 0.0001). We also found a distinct interaction effect between age and intelligibility (F(1, 49) = 21.536, *p* = 1.1e-05, Figure 4D, cingulum). The Bonferroni corrected post-hoc test also revealed a significant difference between the forward and backward conditions in the young group (*p* = 1.1e-07), but not in the older group (*p* = 1.000). Additionally, we found a significant difference between the forward condition in the young group and the forward condition in the older group (*p* = 0.0005), strengthening our observation that the young group is able to distinguish more faithfully between forward and backward speech than the older group. In the cingulate cortex, we also found a significant difference between the forward condition in the young group and the backward condition in the older group (p = 0.002). Here, we observe an additional effect in the older group, conveying that the ageing brain fails to distinguish between intelligible and unintelligible speech, and even exhibits a reverse pattern by tracking the presented backward speech more than the young group.

**Figure 4:**
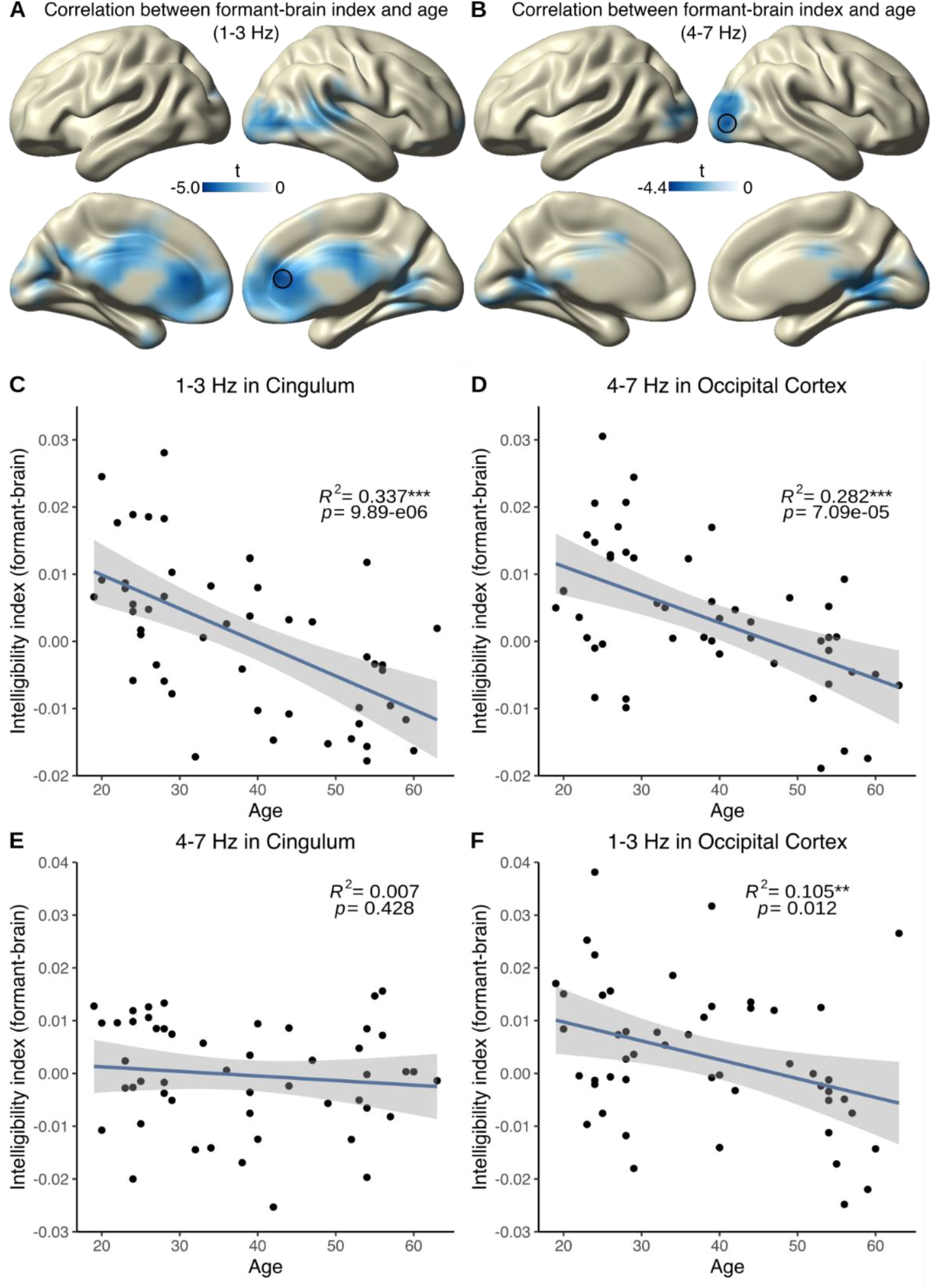
Statistical values of the voxelwise correlation of the formant-brain index with age split between delta-band and theta-band. A) Tracking of the intelligibility index in the delta-band (1-3 Hz, p < 0.05, cluster-corrected) indicates a strong decrease of intelligibility tracking in cingulate cortex and frontal areas. Black circle indicates lowest t-value extracted for C and F. B) Tracking of the intelligibility index in the theta-band (4-7 Hz, p < 0.05, cluster-corrected) indicates a strong decrease of intelligibility tracking in visual areas. Black circle indicates lowest t-value extracted for D and E. C) Correlation of formant-brain intelligibility index in the voxel with the lowest t-value extracted from A (cingulate cortex) showing a significant decrease with age (p = 9.885e-06) in the delta-band. D) Correlation of formant-brain intelligibility index in the voxel with the lowest t-value extracted from B (visual cortex) showing a significant decrease with age (p = 7.089e-05) in the theta-band. E) Correlation of formant-brain intelligibility index in the voxel with the lowest t-value extracted from A (cingulate cortex) showing no significant decrease with age (p = 0.428) in the theta-band. F) Correlation of formant-brain intelligibility index in the voxel with the lowest t-value extracted from B (occipital cortex) showing a significant decrease with age (p = 0.012) also in the delta-band. Blue lines depict regression lines, shaded areas depict standard error of mean (SE).

### 3.5 Different frequency-bands show an age-related decline in different brain regions

As a last step, we investigated if different frequency bands are impacted differently by age-related decline. Therefore, we repeated the analysis steps explained in 3.3, meaning that we calculated again a voxelwise correlation between the intelligibility index separately for our coherence conditions (lip-brain, envelope-brain, formant-brain and pitch-brain) and the age of the participants, but this time separately for the delta-band (1-3 Hz) and the theta-band (4-7 Hz). For the delta-band, we again found a significant correlation between age and the intelligibility index just for the formant-brain index (*p* = 0.002, cluster-corrected). This effect was strongest in cingulate cortex (lowest *t*-value: −4.991, MNI [0 40 10], Figure 4A). No correlation occurred between age and the other indices (lip-brain index: *p* = 0.833; envelope-brain index: *p* = 0.268; pitch-brain index: *p* = 0.166, all cluster-corrected). Repeating the analysis for the theta-band revealed a similar picture: While we could find a significant correlation between the formant-brain index and age (*p* = 0.018, cluster-corrected) which was strongest in visual cortex (lowest t-value: −4.394, MNI [40 −90 0], Figure 4B), we did not find it for the remaining indices and age (lip-brain index: *p = 1*; envelope-brain index: *p* = 0.096; pitch-brain index: *p* = 0.675, all cluster-corrected). These results display a differential spatial pattern for different frequency bands: While tracking of intelligible speech in the theta-band declines reliably in visual cortex, tracking of the slower delta-band rather declines in cingulate cortex and frontal areas. We then extracted the values from the voxel with the lowest *t*-value (i.e. the most significant negative one) respectively for both frequency bands (delta-band: cingulate cortex, MNI [0 40 10]; theta-band: visual cortex, MNI [40 −90 0]) and again fitted a linear model for the formant-brain index to further clarify the effects found in the whole brain analysis. Age could significantly predict the formant-brain index in the delta-band in cingulate cortex (F(1, 48) = 24.4, *p =* 9.885e-06, *η^2^* = 0.337, Figure 4C) and in the theta-band in visual cortex (F(1, 48) = 18.92, *p* = 7.089e-05, *η^2^* = 0.282, Figure 4D). To further clarify if the tested relationship is specific to a certain frequency band and brain region, we also tested the vice versa relationship (i.e. the relationship between age and theta-band in cingulate cortex and the relationship between age and delta-band in occipital cortex). We found that while age could not significantly predict the formant-brain index in the theta-band in cingulate cortex (F(1, 48) = 0.637, *p* = 0.429, *η^2^* = 0.01, Figure 4E), it could significantly predict the formant-brain index in the delta-band in occipital cortex (F(1, 48) = 6.757, *p* = 0.012, *η^2^* = 0.123, Figure 4F). This suggests that while the ability of the cingulate cortex to transform visual into phonological information declines just in the delta-band, the occipital cortex shows a decline over a broad range of frequencies and therefore in general visual speech processing.

## 4 Discussion

Our study illustrates that during lip reading, the visual cortex represents multiple features of the speech signal in low frequency bands (1-7 Hz), importantly including the corresponding (unheard) acoustic signal. It has previously been shown that the visual cortex is able to track the intelligible global envelope (unheard acoustic speech envelope; Hauswald et al. 2018). We demonstrate here that the visual cortex is also able to track the modulation of intelligible spectral fine-details (unheard formants and pitch). Furthermore, we found that ageing is associated with a deterioration of this ability not only in the visual cortex, but also in the cingulate cortex. Disentangling delta and theta-band revealed that while the age-related decline of formant tracking is independent of frequency bands in visual cortex, it is unique in cingulate cortex for the delta-band. Our results suggest that visuo-phonological transformation processes are sensitive to age-related decline, in particular with regards to the modulation of unheard spectral fine-details.

### Visuo-phonological transformation processes are observable for global amplitude modulations and spectral-fine detail modulations

As expected, the current study replicates the main finding from Hauswald et al. (2018) showing a visuo-phonological transformation process in visual cortex for the unheard speech envelope in an Italian speaking sample. Our study using a German speaking sample suggests that the postulated visuo-phonological transformation process at the level of the visual cortex is generalizable across languages. This is unsurprising as it is in line with studies on the speech envelope spectrum which show robust amplitude peaks between 3.5 and 4.5 Hz regardless of language (Poeppel & Assaneo, 2020), providing evidence that different languages carry similar temporal regularities not only for auditory properties, but also for visual properties (Chandrasekaran et al., 2009). We argue that this similarity is a key property for making the postulated visuo-phonological transformation process transferable to other languages.

By investigating different properties of auditory speech (global modulations vs. fine-detailed modulations) and how they are tracked by the human brain, our results are furthermore adding an important part to the understanding of how visual speech contributes to speech processing in general. As lip movements and amplitude modulations are highly correlated (Chandrasekaran et al., 2009), it is highly probable that amplitude modulations can be inferred by lip movements alone as a learned association. Here we can show that the brain is also able to perform a more fine-coarsed tracking than initially thought by especially processing the spectral fine-details that are modulated near the lips, another potentially learned association between lip-near auditory cues (i.e. merged F2 and F3 formants) and lip movements (Plass et al., 2020). Additionally, it is not only formants that are subject to visuo-phonological transformation, but also the fundamental frequency, as seen in our results. This is in line with a recent study which shows that closing the lips is correlated with the tone falling (Garg et al., 2019). How those modulations are influenced by behavioural measures still needs to be discussed. Some studies suggest that enhanced lip reading abilities go in line with higher activation in visual areas in persons with a cochlear implant (e.g. Giraud et al., 2001). Our present results do not suggest that strong visuo-phonological transformation processes are sufficient for improved lip reading abilities. Yet, they may be most useful in disambiguating auditory signals in difficult listening situations.

### Tracking of unheard formants accompanying lip movements is mostly affected in ageing

With regards to the ageing effect, we could show that various neural tracking mechanisms are differentially affected. Our study presents that tracking of unheard formants, especially the combined F2 and F3 formants, is declining with age, while there is still a preserved tracking of purely visual information (as seen in the lip-brain index, Figures 3C and 3F). Meanwhile, the tracking of the unheard speech envelope and pitch signify an inconclusive picture: While tracking of those properties seem to be preserved to some extent, both are showing a tendency to diminish with age.

Especially the formants and the pitch are part of the temporal fine-structure (TFS) of speech and are crucial for speech segregation or perceptual grouping for optimal speech processing in complex situations (Alain et al., 2017; Bregman et al., 1990). The TFS is different from the acoustic envelope in a sense that it does not display “coarse” amplitude modulations of the audio signal but rather fluctuations that are close to the centre frequency of certain frequency bands (Lorenzi et al., 2006). Hearing-impaired older participants show a relative deficit of the representation of the TFS compared to the acoustic envelope (Anderson et al., 2013; Lorenzi et al., 2006). The TFS also yields information when trying to interpret speech in fluctuating background noise (Moore, 2008). Other studies also point to the fact that especially when having cochlear hearing loss along with a normal audiometric threshold, the interpretation of the TFS is reduced, resulting in diminished speech perception under noisy conditions (Lorenzi et al., 2009). This suggests that hearing-impaired subjects mainly seem to rely on the temporal envelope to interpret auditory information (Moore & Moore, 2003), while normal hearing subjects can also use the presented temporal fine-structure. Interestingly, we found that even when the TFS is inferred from lip movements, there is a decline in the processing of spectral fine-details with age independent of hearing loss. Our results suggest that the visuo-phonological transformation of certain spectral fine-details like the formants are impacted the most in ageing, whereas the transformation of the pitch (or fundamental frequency) reveals a more complex picture: We find preserved tracking of the unheard pitch contour in occipital cortex, but a decline with age in the cingulate cortex. Interestingly, the cingulate cortex has been found to show higher activation as response to processing of degraded speech (Erb & Obleser, 2013), pointing to a possible compensatory mechanism when processing distorted speech. How this altered processing of the unheard pitch (or fundamental frequency) accompanying lip movements in cingulate cortex has an impact on speech understanding needs to be discussed in further studies.

Further investigating the effects shown in our correlational analysis revealed that older participants seem to be less able to distinguish between forward and backward unheard speech (unheard formants) and that younger individuals show enhanced tracking of intelligible speech (Figure 3D). This could point to the fact that the older population is losing the gain of differentiating intelligible from unintelligible speech, obviously resulting in a less successful visuo-phonological transformation process. Other studies suggest that the older population seems to inefficiently use their cognitive resources, showing less deterioration of cortical responses (measured by the envelope reconstruction accuracy) to a foreign language compared to younger individuals (Presacco et al., 2016b) and also an association between cognitive decline and increased cortical envelope tracking or even higher synchronization of theta (Goossens et al., 2016). Auditory processing is also affected both in midbrain and cortex in age, exhibiting a large reduction of speech envelope encoding when presented with a competing talker, but at the same time a cortical overrepresentation of speech regardless of the presented noise, suggesting an imbalance between inhibition and excitation in the human brain (Presacco et al., 2016a) when processing speech. Other studies add to this hypothesis by showing decreasing alpha modulation in the ageing population (Henry et al., 2017; Vaden et al., 2012), strengthening the assumption that there is an altered interaction between age and cortical tracking even in the visual modality that needs to be investigated further.

Considering all acoustic details accompanying lip movements we still see a tendency of the speech envelope tracking to decline with age, suggesting that the transformation of the global speech dynamics could also be deteriorating. Overall, our results provide evidence that the transformation of fine-grained acoustic details seem to decline more reliably with age, while the transformation of global information (in our case the speech envelope) seems to be less impaired.

### Possible implications for speech processing in challenging situations

Our findings raise the question of how the decline in processing of unheard spectral fine-details negatively influences other relevant aspects of hearing. In light of aforementioned studies from the auditory domain of speech processing, we propose some thoughts on the multi-sensory nature of speech and how different sensory modalities can contribute to speech processing abilities under disadvantageous conditions (both intrapersonal and environmental).

As mentioned in the previous section, optimal hearing requires processing of both the temporal fine structure and the global acoustic envelope. However, especially under noisy conditions, processing the TFS becomes increasingly important for understanding speech. Ageing in general goes along with reduced processing of the TFS (Anderson & Karawani, 2020) and this deteriorating effect seems to be even more detrimental when ageing is accompanied by hearing loss (Anderson et al., 2013). Since listening in natural situations usually is a multi-sensory (audiovisual) phenomenon, we argue that the impaired visuo-phonological transformation process of the TFS adds to the difficulties of older (also audiometrically normal hearing) individuals to follow speech in challenging situations. To follow up this idea, future studies will need to quantify the benefit of audiovisual versus (unimodal) auditory processing, depending on different visuo-phonological transformation abilities.

Our results also have implications for listening situations when relevant visual input from the mouth area is obscured, a topic which has gained enormously in significance due to the wide adoption of face masks to counteract the spread of SARS-CoV-2. In general, listening becomes more difficult and performance declines when the mouth area is obscured (Brown et al., 2021; Giovanelli et al., 2021). While face masks may diminish attentional focusing as well as temporal cues, our work suggests that they also deprive the brain of deriving the acoustic TFS from the lip movements especially in the formant frequency range which are modulated near the lips (F2 and F3). This issue, which should become relevant particularly in noisy situations, may be aggravated by the fact that face masks (especially highly protective ones) impact sound propagation of frequencies between 1600-6000 Hz with a peak around 2000 Hz (Caniato et al., 2021). Thus, face masks diminish relevant formant information in both sensory modalities. This could disproportionately affect hearing impaired listeners, an urgent question that should be followed up by future studies.

Overall, considering both the auditory and visual domain of speech properties, we suggest that the underlying cause of speech processing difficulties in naturalistic settings accompanying age or hearing impairment is more diverse than previously thought. The visual system provides the proposed visuo-phonological transformation process as an important mechanism for optimal speech understanding and crucially supports acoustic speech processing.

### Occipital cortex and cingulate cortex show different tracking properties dependent on the frequency-band

With regards to different frequency bands, our results could yield important insights into different brain regions showing distinct formant tracking properties: While we find a robust decline of delta-band tracking with age in both occipital and cingulate cortex, theta-band tracking is reliably declining only in occipital areas. In general, theta is corresponding to the frequency of syllables and to the modulations in the amplitude envelope (Gross et al., 2013; Keitel et al., 2018; Meyer, 2018; Poeppel & Assaneo, 2020), whereas delta seems to process phrasal chunks based on acoustic cues (Ghitza, 2017; Keitel et al., 2018) and is therefore responsible for a general perceptual chunking mechanism (Boucher et al., 2019). Our results also show that the visual cortex extracts information provided by the perception of the lip movements and connects them with phonological information that is already learned. This points to a possible top-down influence of stored syntactic information provided by delta-band tracking, which also seems to be deteriorating with increasing age both in occipital and cingulate cortex. Interestingly, age-related hearing loss also leads to a volume reduction in anterior cingulate cortex (Slade et al., 2020), which in turn also leads to more memory impairments and cognitive deficits (Belkhiria et al., 2019). These and our current results strengthen the notion that the cingulate cortex has an important function also in visual speech processing, as this also goes in line with the mentioned compensatory mechanism in anterior cingulate cortex (ACC) (Erb & Obleser, 2013). Together with the findings of the current study, this involvement of the cingulate cortex in speech processing (or in general the cingulo-opercular network; Peelle (2018)) underlines the fact that there seems to be a maladaptive processing strategy in frontal areas. To fully understand the mechanisms behind this visuo-phonological transformation process without the influence of ageing in distinct brain regions and frequency bands, it would be advisable for future studies to focus on younger individuals, especially since this study is the first to investigate the tracking of spectral fine-details extracted from the spectrogram.

## 5 Conclusion

The current study demonstrates that the visual cortex is able to track intelligible unheard spectral-fine detailed information just by observing lip movements. Crucially, we present a differential pattern for the processing of global (i.e. envelope) and spectral fine-detailed intelligible information, with ageing affecting in particular tracking of spectral speech information (or the TFS), while showing partly preserved tracking of global modulations. Furthermore, we see a distinct age-related decline of tracking dependent on the brain region (i.e. visual and cingulate cortex) and on the frequency-band (i.e. delta and theta-band). The results presented here may have important implications for hearing in the ageing population, suggesting that hearing difficulties could also be exacerbated in natural audiovisual environments as a result of reduced capacities of visual benefit. With respect to the current pandemic situation, our results can provide a novel important insight on how missing visual input (e.g. when carrying face masks) is critical for speech comprehension.

## 7 Acknowledgements

This work is supported by the Austrian Science Fund, P31230 (“Audiovisual speech entrainment in deafness”).

## 8 Pre-registration

The first part of the study analyses was pre-registered prior to the research being conducted under https://osf.io/ndvf6/.

## 9 Data availability

The “mat” and “csv” files containing the data shown in the figures, along with the MATLAB code and the R code to recreate the plots, are available under https://osf.io/ndvf6/. Readers seeking access to the original, non-resampled data (~430 GB) should contact the lead author. Access will be granted in accordance with ethical procedures governing the reuse of sensitive data.

